# Genome-wide analysis of *Enterococcus faecalis* genes that facilitate interspecies competition with *Lactobacillus crispatus*

**DOI:** 10.1101/2024.10.14.618276

**Authors:** Ling Ning Lam, Kathryn E. Savage, Camille N. Shakir, José A. Lemos

**Affiliations:** Department of Oral Biology, College of Dentistry, University of Florida, Gainesville, FL, USA

**Keywords:** enterococci, lactobacilli, *E. faecalis*, interspecies competition, bacterial antagonism

## Abstract

Enterococci are opportunistic pathogens notorious for causing a variety of infections. While both *Enterococcus faecalis* and *Lactobacillus crispatus* are commensal residents of the vaginal tract, the molecular mechanisms that enable *E. faecalis* to outcompete *L. crispatus,* and consequently cause vaginal infections remains unknown. To begin to address this, we need to gain a better understanding of the competitive interactions between *E. faecalis* and *L. crispatus*. Here, we show that *L. crispatus* eradicates *E. faecalis* in a contact-independent manner. Using transposon sequencing to identify *E. faecalis* OG1RF transposon (Tn) mutants that are either under-represented or over-represented when co-cultured with *L. crispatus*, we found that Tn mutants with disruption in the *dltABCD* operon, that encodes the biosynthetic pathway for D-alanylation of teichoic acids, and *OG1RF_11697* encoding for an uncharacterized hypothetical protein are more susceptible to killing by *L. crispatus*. Inversely, Tn mutants with disruption in *ldh1,* that encodes for L-lactate dehydrogenase, are more resistant to *L. crispatu*s killing. Using the *Galleria mellonella* infection model, we show that co-injection of *L. crispatus* with *E. faecalis* OG1RF enhances larvae survival while this *L. crispatus*-mediated protection was lost in larvae co-infected with either *L. crispatus* and *E. faecalis* Δ*ldh1* or Δ*ldh1*Δ*ldh2* strains. Last, using RNA sequencing to identify *E. faecalis* genes that are differently expressed in the presence of *L. crispatus*, we found major changes in expression of genes associated with glycerophospholipid metabolism, central metabolism and general stress responses. The findings in this study provide insights on how *E. faecalis* mitigate assaults by *L. crispatus*.

**IMPORTANCE:** *Enterococcus faecalis* is an opportunistic pathogen notorious for causing a multitude of infections. While *E. faecalis* must compete with *Lactobacillus crispatus* to cause vaginal infections, how *E. faecalis* overcomes *L. crispatus* killing remains unknown. We show that *L. crispatus* eradicates *E. faecalis* temporally in a contact-independent manner. Using high throughput molecular approaches, we identified genetic determinants that enable *E. faecalis* to compete with *L. crispatus*. This study represents an important first step for the identification of adaptive genetic traits required for enterococci to compete with lactobacilli.

## INTRODUCTION

Enterococci, residents of the gastrointestinal and urogenital tracts, are notorious for causing a variety of opportunistic infections including but not limited to catheter-associated urinary tract infections (CAUTI) ^1,2^, antibiotic-induced intestinal dysbiosis ^3,4^, surgical site and diabetes-associated wound infections ^5–7^, endodontic infections ^8,9^, and infective endocarditis (IE) ^10–12^. Notably, enterococci are amongst the top five bacterial pathogens associated with bloodstream infections, surgical site infections, and CAUTI ^13,14^, and is the third most common bacterial agent of IE ^15^. Armed with the ability to acquire antibiotic resistance ^16^, and a fastidious nature ^17^, enterococcal infections became even more challenging to treat when these microbes are found in polymicrobial biofilm communities ^18,19^. In addition, these polymicrobial biofilms can also serve as a nidus for systemic dissemination and a reservoir for horizontal transfer of resistance genes ^18,20^. The robust nature of biofilms has posed major challenges in industrial, agricultural, and clinical settings ^21–23^. For these reasons, there has been a mounting interest to understand the molecular processes driving species cooperation and competition within complex biofilm communities to inform tactics for clinical interventions ^24^.

While not an exhausting list, previous studies have investigated microbial interactions of enterococci ^18^. In one study, it was reported that *Enterococcus faecalis*, the most prevalent enterococci in humans, suppresses *Clostridium perfringens* growth through production of a bacteriocin ^25^. Around the same time, different groups showed that *E. faecalis* can inhibit production of the botulinum neurotoxin by *Clostridium botulinum* ^26^, biofilm formation by *Streptococcus mutans* ^27^, and virulence of the *Candida albicans* yeast in a biofilm-associated oropharyngeal candidiasis mouse model ^28,29^. Other studies have showed that co-culturing of *E. faecalis* with *Escherichia coli* results in suppression of pro-inflammatory responses ^30^, augmentation of the dual-species biofilm biomass ^31^, and heighten *E. coli* burden in a co-infection mouse incisional wound model ^32^. Another major opportunistic pathogen of infected wounds, *Pseudomonas aeruginosa* is frequently co-isolated with *E. faecalis* in polymicrobial wound infections ^22,33^. Like *E. coli*, co-culturing of *E. faecalis* and *P. aeruginosa* augmented biofilm biomass due to Psl- and Pel-mediated matrix polysaccharide production by *P. aeruginosa* ^34,35^. Paradoxically, a recent study indicated that *E. faecalis* inhibits *P. aeruginosa* growth by lowering environmental pH and L-lactate-mediated iron chelation ^36^. In an antibiotic perturbed gastrointestinal mouse model, co-infection of *E. faecalis* with *C. difficile* enabled sharing of fermentable amino acids, leucine and ornithine synthesized by *E. faecalis* while depletion of environmental arginine by *E. faecalis* acts as a metabolic cue to trigger upregulation of virulence in *C. difficile* ^37^. Finally, a recent study showed that exogenous heme released from the growth of *Staphylococcus aureus* augment *E. faecalis* biofilm, and this is facilitated by *E. faecalis* gelatinase activity that aids in the extraction of heme from *S. aureus* hemoproteins ^38^. Together, these studies reveal that specific bacterial pairs and the environmental context dictate whether *E. faecalis* interactions will be antagonistic or mutualistic.

Lactobacilli have also been shown to antagonize a variety of microbial pathogens^39–41^. Due to their beneficial role in urogenital and gastrointestinal microbiome homeostasis, several members of this genus have been used as probiotics ^42,43^. Of note, lactobacilli (typically *L. crispatus, L. iners, L. gasseri*, or *L. jensenii*) ^44–46^ constitute more than 50% of the microbial population of the vaginal tract and their abundance is strongly associated with vaginal health ^45,47^. In fact, depletion of lactobacilli in the vaginal flora is a clear sign of vaginal dysbiosis and, consequently, disease onset ^48^. The robust antagonistic capabilities of lactobacilli enable these commensal organisms to curb outgrowth of potential vaginal pathogens and maintain vaginal eubiosis ^49,50^.

In recent years, *E. faecalis* has been implicated as an emerging vaginal pathogen ^51–53^, with separate reports documenting its outgrowth and prevalence in vaginal swabs of patients suffering from bacterial vaginosis, and more recently, aerobic vaginitis ^54–56^. Apart from a study implicating ethanolamine catabolism and the type VII secretion system in *E. faecalis* colonization of the vaginal tract ^57^, our understanding of the pathophysiology of *E. faecalis* in this microenvironment remains incomplete. Perturbation of vaginal microbiome homeostasis ^58,59^ and the depletion of vaginal lactobacilli population ^60^ often precede onset of vaginal infection. Concurring with this notion, a recent study by France and colleagues reported that vaginal microbial communities with overgrowth of vaginal pathogens (CST IV-C1, *Streptococcus* dominated; CST IV-C2, *Enterococcus* dominated; CST IV-C3, *Bifidobacterium* dominated; CST IV-C4, *Staphylococcus* dominated) have drastically reduced abundance of vaginal lactobacilli ^45^. By understanding the competitive interactions between *E. faecalis* and *L. crispatus*, a biomarker of vaginal health^61^, we could then find explanations for *E. faecalis* ability to outcompete vaginal lactobacilli and persist in the vaginal tract. Using *in vitro* dual-species biofilm models, we show that *L. crispatus* eradicates *E. faecalis* temporally in a contact-independent manner. In addition, we showed that *E. faecalis*-*L. crispatus* co-infection protects the *Galleria mellonella* larvae against *E. faecalis*-mediated killing. Using transposon sequencing (Tn-seq) and RNA sequencing (RNA-seq) approaches, we identified specific genes and pathways that are negatively or positively associated with the ability of *E. faecalis* to compete with *L. crispatus*. Altogether, this study represents a first step for the identification of adaptive traits that allows *E. faecalis* to compete with lactobacilli, providing insights into the mechanisms that enable the persistence of *E. faecalis* in the vaginal tract.

## RESULTS

### *L. crispatus* eradicates *E. faecalis* in a temporal manner

Because the prevalence of *L. crispatus* is an indicator of a healthy vaginal tract^44,61^, and *E. faecalis* is an understudied bacterial agent of vaginal infection ^51–53^, we sought to investigate the microbial interactions between these two species. In the first series of experiments, we determined *E. faecalis* OG1RF and *L. crispatus* VPI3199 growth and long-term viability as either single-species or mixed-species macro-colony biofilms (**Fig 1A**). While *E. faecalis* alone grew to higher cell density as compared to *L. crispatus*, the viability of *L. crispatus,* alone or mixed with *E. faecalis,* remained unchanged (**Fig 1B**). However, there was a stepwise reduction in colony-forming units (CFUs) of *E. faecalis* when co-cultured with *L. crispatus* for 24 hrs (∼0.5-log reduction), 72 hrs (2-log reduction) and 120 hrs (∼2.5-log reduction) that was not observed in the single-species *E. faecalis* macro-colony control (**Fig 1B**). These results indicate that *L. crispatus* kills *E. faecalis* over time. To determine if *L. crispatus-*mediated killing was conserved at the genus level, we probed the killing capacity of *L. crispatus* VPI 3199 against a panel of *E. faecalis* and *E. faecium* clinical and non-clinical strains. Like *E. faecalis* OG1RF, all other enterococcal strains lost viability when co-cultured with *L. crispatus,* with the vancomycin-resistant *E. faecalis* V583 showing the greatest rate of killing (∼4-log after 72 hrs) (**Fig 1C**). In addition, five different clinical strains of *L. crispatus* also showed equivalent capacities to kill *E. faecalis* OG1RF when mixed (**Fig 1D, Fig S1**). Lastly, we showed that other lactobacilli, *L. casei* and *L. rhamnosus*, can efficiently kill *E. faecalis* over time (**Fig S2**). Besides macro-colony biofilms, we also grew *E. faecalis* OG1RF and *L. crispatus* VPI3199 biofilms statically, either single or mixed species, using tissue culture plates for up to 72 hrs. In this experimental set up, *L. crispatus* VPI completely eradicated *E. faecalis* OG1RF after 72 hrs. Again, the viability of *L. crispatus* did not differ (**Fig S3**). Collectively, these results provide compelling evidence that *L. crispatus* and other lactobacilli can efficiently kill *E. faecalis* in dual-species biofilms *in vitro*.

**Fig 1.**
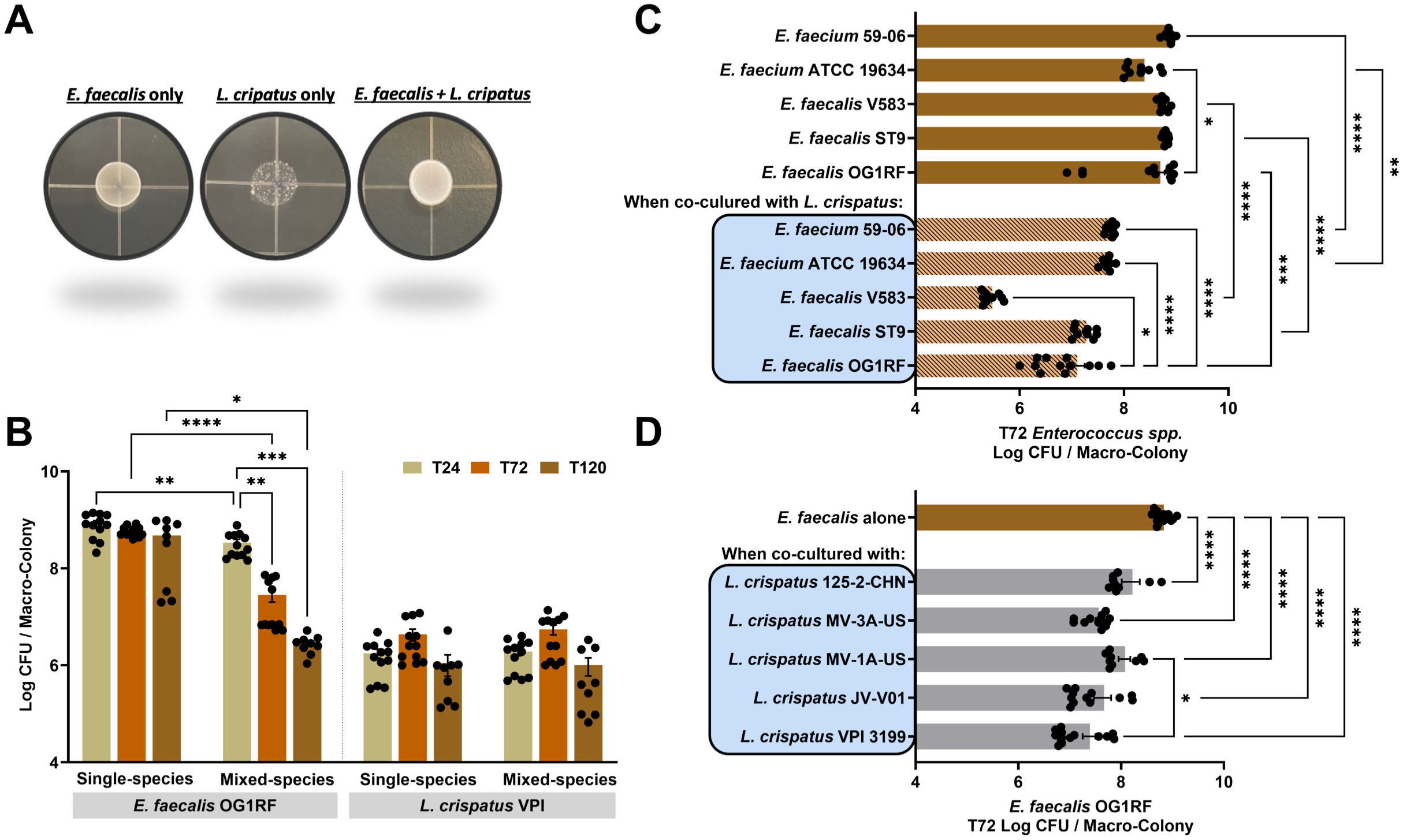
*L. crispatus* antagonizes growth of enterococci. (A) Representative image of macro-colony biofilms on a MRS agar plate. (B) Colony-forming units (CFU) recovered from *E. faecalis* OG1RF and *L. crispatus* VPI 3199 grown respectively either as single-species macro-colony biofilm, or as dual-species (*E. faecalis* + *L. crispatus*) macro-colony biofilm for 24 hrs, 72 hrs and 120 hrs post incubation. (C) Colony-forming units (CFU) recovered from selected *E. faecalis* and *E. faecium* strains respectively grown either as single-species macro-colony biofilm, or grown as dual-species macro-colony biofilms after 72 hrs incubation. (D) Colony-forming units (CFU) recovered from *E. faecalis* OG1RF grown either as single-species macro-colony biofilm or grown respectively with selected *L. crispatus* strains as dual-species macro-colony biofilm after 72 hrs incubation. For B-F, data points represent the average of 9-12 biological replicates, collated from at least three repeated experiments. Statistical analysis was performed using Brown-Forsythe ANOVA test with Welch’s correction. Error bars represent the standard error of margin (SEM). * *p* ≤ 0.05, ** *p* ≤ 0.01, *** *p* ≤ 0.001, **** *p* ≤ 0.0001.

### Mechanism of killing is contact-independent

To obtain insights into the mechanism(s) of *L. crispatus*-mediated killing of *E. faecalis*, we performed a series of *in vitro* static biofilm experiments using the 72 hr time point (∼7-log killing, refer to **Fig S3**) as the read-out. To probe whether killing is contact-dependent, we grew single-species *E. faecalis* and *L. crispatus* biofilms together in a tissue culture plate, separated by a Transwell™ permeable membrane insert that prevent physical contact while allowing exchange of metabolites and other types of molecules in the same well. Because CFUs recovered from *E. faecalis* biofilms seeded on the surface of the Transwell™ membrane insert was drastically reduced (∼7-log; similar levels to those observed in **Fig S3**), while CFUs of *L. crispatus* biofilms at the bottom of the culture plate remained unchanged (**Fig 2A**), we concluded that secreted molecule(s) are responsible for the eradication of *E. faecalis*. Next, we switch to a plate-based macro-colony biofilm antagonism assay to investigate if *L. crispatus* secreted, diffusible molecule(s) are produced in the absence of *E. faecalis*. While simultaneous inoculation of *E. faecalis* and *L. crispatus* did not result in a zone of clearing at T0 and minimally affected *E. faecalis* growth when inoculated 24 hr later (T24), spotting of *E. faecalis* macro-colony biofilms 48 and 72 hrs after the growth of *L. crispatus* macro-colony biofilms resulted in clear zones of inhibition (**Fig 2B**). This result indicates that *L. crispatus* mediated growth inhibition of *E. faecalis* occurs at later time points and is not associated with the presence of *E. faecalis*. While *L. crispatus* bacteriocins have not been described, previous studies have identified phenyl-lactic acid (PLA), present in cell-free supernatants of *L. crispatus*, as a bactericidal compound with wide spectrum activity against other bacteria, including *E. faecalis* ^62^. PLA was shown to cause cell membrane damage providing, at least in part, an explanation for its antimicrobial activity ^63^. To test if PLA produced by lactobacilli can be accounted for the killing of *E. faecalis* observed here, we determined the inhibitory concentration of PLA against *E. faecalis* OG1RF or *L. crispatus* VPI. Based on the observation that similar concentrations of PLA were inhibitory to both species (**Fig S4**), we conclude that it is unlikely that *L. crispatus* secreted PLA alone is responsible for the killing of *E. faecalis*. Moreover, enterococci also secrete PLA, and the extracellular concentrations of PLA produced by either lactobacilli or enterococci that has been reported ^64,65^, approximately 1 mM (0.16 mg/mL), is not inhibitory to *E. faecalis*.

**Fig 2.**
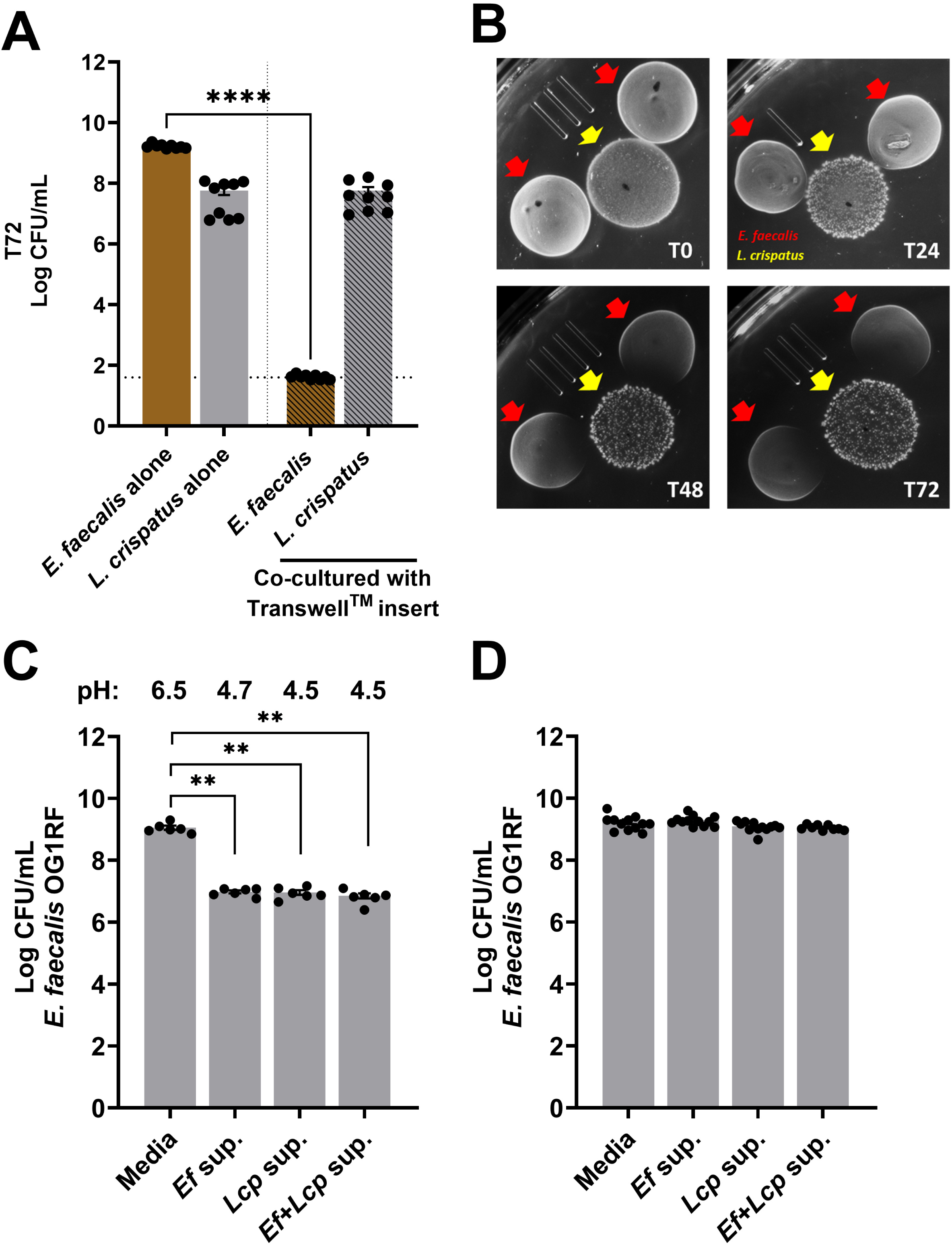
*L. crispatus* antagonism of *E. faecalis* is facilitated in a contact-independent manner. (A) Colony-forming units (CFU) recovered from *E. faecalis* OG1RF and *L. crispatus* VPI 3199 grown statically for 72 hrs in MRS media as single-species biofilm, either in separate wells or in the same well separated by a Transwell™ membrane insert that prevents physical contact between cells. *L. crispatus* biofilm is grown on the flat surface of the well in the tissue culture plate, whereas *E. faecalis* biofilm is seeded on the surface of the Transwell^TM^ membrane insert. Dotted line indicates limit of detection (LOD), CFU<42.5. (B) Representative images of spot antagonism assay showing growth inhibition of *E. faecalis* when *L. crispatus* macrocolony biofilms were established at the same time (T0), 24 hr, 48 hr or 72 hr before inoculating *E. faecalis*. Colony-forming units (CFU) recovered from *E. faecalis* OG1RF growth after 24 hrs in MRS media mixed with 72 hrs cell-free biofilm supernatant isolated from single-species and dual-species biofilms at equal ratio, either (C) pH-unadjusted or (D) adjusted to pH 6.5 to mirror the MRS media. For A, C and D, data points represent an average of 9-12 biological replicates, collated from at least three repeated experiments. Statistical analysis was performed using Brown-Forsythe ANOVA test with Welch’s correction. Error bars represent the standard error of margin (SEM). ** *p* ≤ 0.01, **** *p* ≤ 0.0001.

Because *E. faecalis* viability is drastically reduced after 72 hrs post incubation in co-culture (**Fig 2A**), we reasoned that the *L. crispatus* anti-enterococcal diffusible molecule(s) might be of most abundant at this time point. To test this possibility, we mixed cell-free biofilm supernatants collected from either single or mixed species *L. crispatus* biofilms grown for 72 hrs, with fresh MRS in a 50:50 ratio to monitor *E. faecalis* growth and viability. Because of the possibility that acidic pH of the biofilm supernatants may contribute to the killing activity of *L. crispatus* anti-enterococcal diffusible molecule(s), we left the pH unchanged (*Ef* sup, pH 4.7; *Lcp* sup and *Ef+Lcp* sup, pH 4.5; refer to **Fig 3A**). While we were surprised that the CFUs recovered from the growth of *E. faecalis* in *L. crispatus* (*Lcp* sup) or mixed-species (*Ef+Lcp* sup) supernatant were similar to CFUs recovered from growth in its own biofilm supernatant, it is not startling that the acidic pH of these cell-free biofilm supernatants impeded the overall growth of *E. faecalis* when compared to the media control (**Fig 2C**). To rule out that the growth impairment due to acidic pH undermined our ability to observe reduction in enterococcal CFUs due to *L. crispatus* anti-enterococcal diffusible molecule(s), we adjusted the pH of the biofilm supernatants to match those of the MRS media (pH 6.5). While the overall growth of *E. faecalis* is no longer impeded, compared to its own pH-adjusted biofilm supernatant (*Ef* sup) and media control, the viability of *E. faecalis* when grown in *L. crispatus* (*Lcp* sup) or mixed-species (*Ef+Lcp* sup) pH-buffered supernatant, again did not differ (**Fig 2D**). We then increased the biofilm supernatant to media ratio (90:10); still, we did not observe any reduction in CFUs of *E. faecalis* (**Fig S5**). Altogether, we show that the mechanism of killing is contact-independent. But under these conditions tested, we are unable to verify the presence of *L. crispatus* secreted anti-enterococcal diffusible molecule(s) in biofilm supernatants.

**Fig 3.**
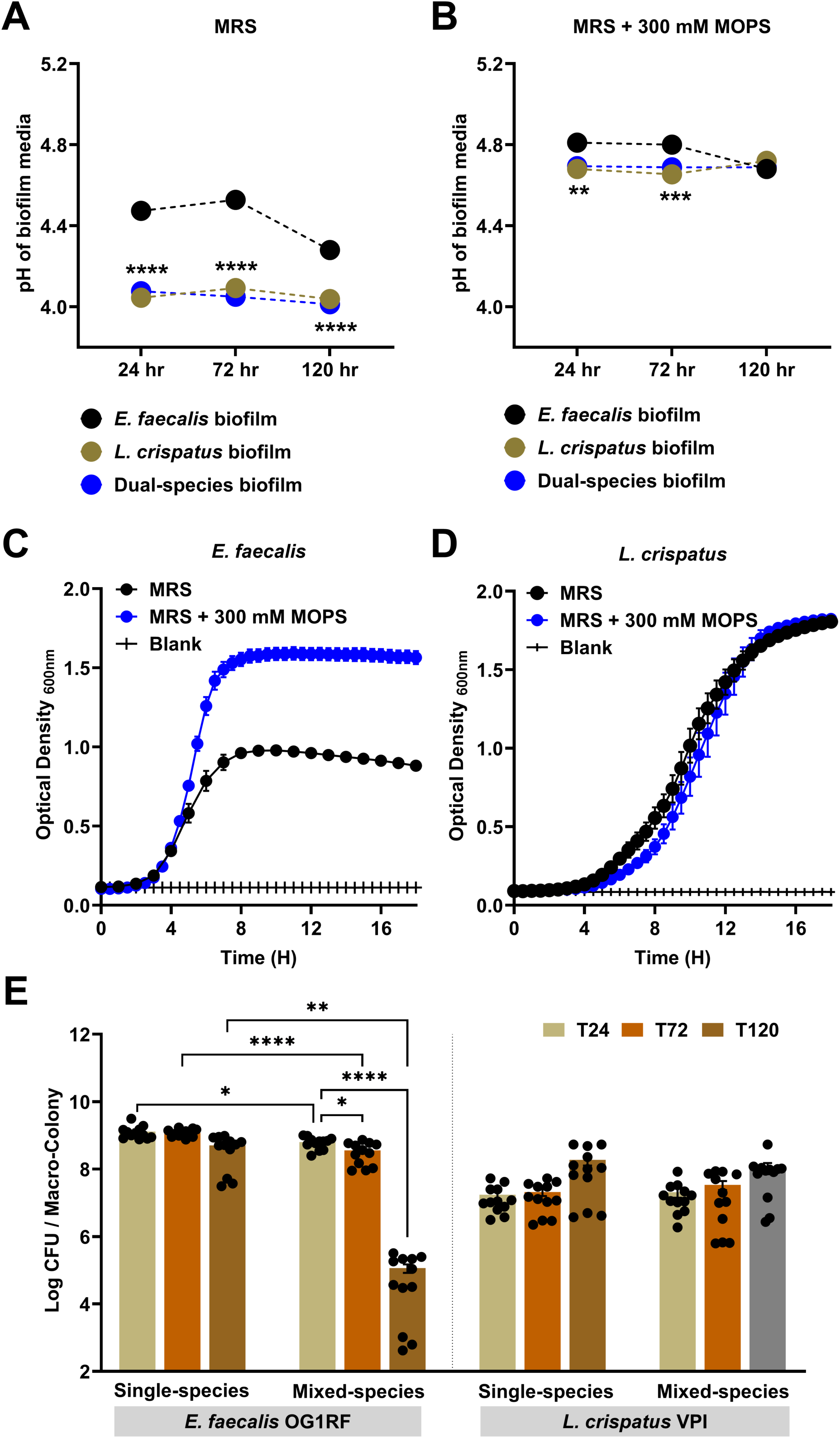
*L. crispatus* antagonistic activity is enhanced in an aciduric environment. pH measurements from *E. faecalis* OG1RF and *L. crispatus* VPI 3199 grown statically between 24-120 hrs either as single-species biofilm, or as dual-species (*E. faecalis* + *L. crispatus*) biofilm in MRS (A) or MRS supplemented with 300 mM MOPS buffer (B). Corresponding growth dynamics of *E. faecalis* (C) and *L. crispatus* (D) in MRS or MRS supplemented with 300 mM MOPS buffer. (E) Colony-forming units (CFU) recovered from *E. faecalis* OG1RF and *L. crispatus* VPI 3199 grown respectively either as single-species macro-colony biofilm, or as dual-species (*E. faecalis* + *L. crispatus*) macro-colony biofilm for 24 hrs, 72 hrs and 120 hrs post incubation in MRS agar supplemented with 300 mM MOPS. For A-E, data points represent an average of 9-12 biological replicates, collated from at least three repeated experiments. For A-B, statistical analysis was performed using two-way ANOVA. For C and D, linear regression of the slope of the exponential growth phase was performed. For E, statistical analysis was performed using Brown-Forsythe ANOVA test with Welch’s correction. Error bars represent the standard error of margin (SEM). * *p* ≤ 0.05, ** *p* ≤ 0.01, *** *p* ≤ 0.001, **** *p* ≤ 0.0001.

### Acidic environment enhances antagonistic activity but is not accountable for the killing of *E. faecalis*

Because lactic acid production by lactobacilli and concomitant environmental acidification has been shown to inhibit growth of other bacteria and is associated with vaginal eubiosis ^49,66^, we next asked whether the lower pH environment created by *L. crispatus* is responsible for the killing of *E. faecalis*. First, we measured the pH of *E. faecalis*, *L. crispatus* and *E. faecalis-L. crispatus* biofilm supernatants at 24, 72 and 120 hr. As expected, *L. crispatus* alone or in co-culture with *E. faecalis* lowered the culture pH (∼4.2) to values below those achieved by *E. faecalis* alone (∼4.3) by 120 hrs post incubation (**Fig 3A**). While we do not anticipate that acidic pH plays a role in the eradication of *E. faecalis* because of two reasons; firstly, killing of *E. faecalis* occurs temporally (**Fig 1B, Fig S3**), and secondly, acidic pH (in static biofilms) remained relatively similar throughout the duration of the experiment (**Fig 3A**), as is expected of these aciduric bacteria, whether alone or mixed. Nonetheless, to rule out that the lower final pH of ∼4.0 driven by *L. crispatus* is implicated in the killing of *E. faecalis* in the mixed-species biofilm, we used 300 mM MOPS (3-(N-morpholino) propanesulfonic acid) to buffer the biofilm pH to around ∼4.6 (*Ef* alone) and ∼4.8 (*Lcp* alone or in co-culture), with all culture conditions reaching the same pH after 120 hr (**Fig 3B**). While the addition of MOPS enhanced the growth of *E. faecalis in vitro,* there was no discernible effect on *L. crispatus* growth (**Fig 3C-D**). As we anticipated, addition of MOPS to MRS agar plates did not rescue *E. faecalis* from eradication in mixed species macro-colony biofilms nor did it affect the viability of *L. crispatus*, albeit reduction in CFUs of *E. faecalis* in co-culture at 72 hrs was less drastic (0.5-log; **Fig 3E**; compared to 2-log reduction in MRS agar, **Fig 1B**). Altogether, these results indicate that acidic environment accelerate but is not the main reason for the killing of *E. faecalis* by *L. crispatus*.

### Identification of transcriptional patterns that enable *E. faecalis* to compete with *L. crispatus*

To obtain insights into how *E. faecalis* responds to *L. crispatus* antagonism, we used RNA sequencing (RNA-seq) to identify transcriptional networks of *E. faecalis* that are altered in *L. crispatus*-*E. faecalis* dual-species macro-colony biofilms. By comparing gene expression profiles of *E. faecalis* grown singly or mixed with *L. crispatus* for 24 hrs, with a cut-off of log_2_FC of ±1 and false discovery rate (FDR) of 0.05, we identified 201 genes down-regulated and 218 genes up-regulated (**Fig 4A, Table S1**). The most up and down -regulated genes, which comprises the top 50 most differentially expressed genes (**Table 1**), according to KEGG pathway analysis (**Fig 4B, Table S2;** *p*-value ≤0.05), were found to be involved in glycerophospholipid metabolism, carbon and sugar utilization, central metabolism, and general stress responses.

**Fig 4.**
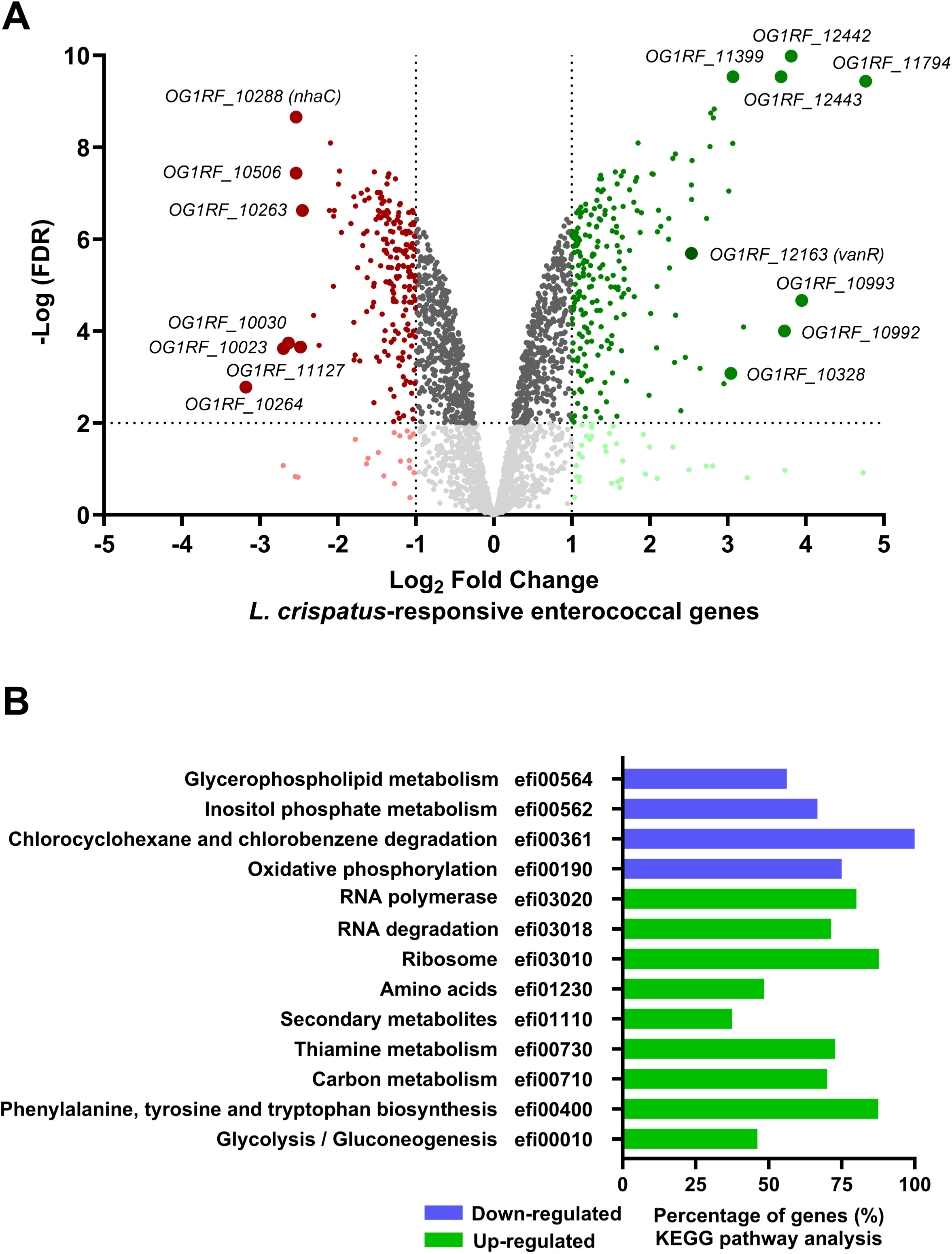
Global transcriptomic changes of *E. faecalis* when co-cultured with *L. crispatus*. (A) Graphical representation of the total number of genes transcriptionally changed relative the degree of log_2_ fold change observed. (B) KEGG pathway analysis of the most significant changes in metabolic pathways. Data shown is averaged from three biological replicates. Refer to methods for information on analysis.

**Table 1.**
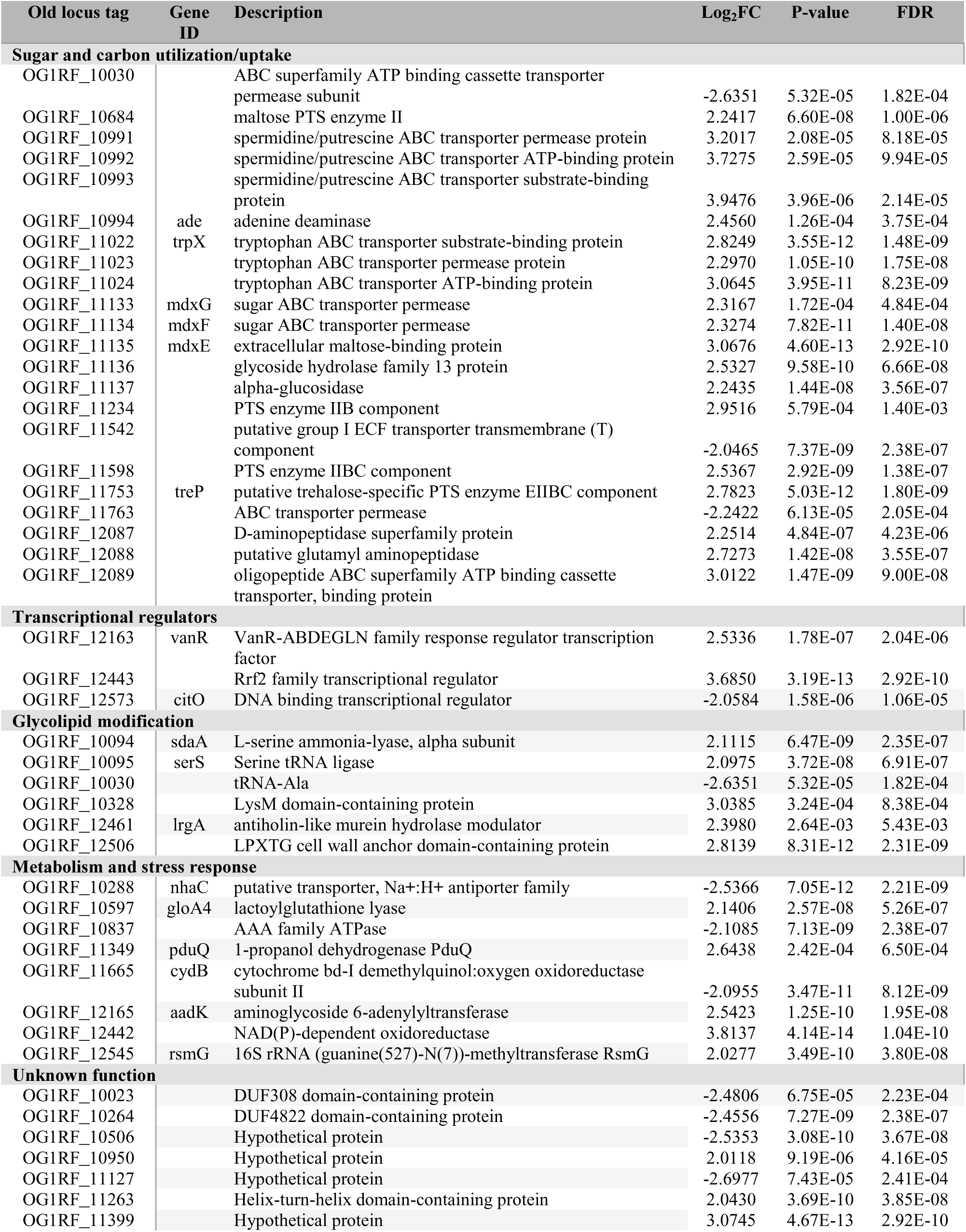
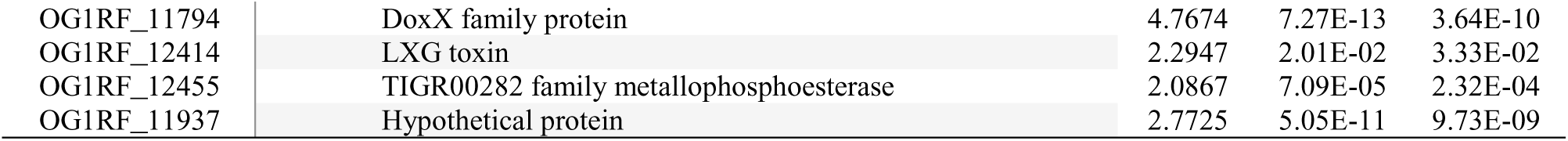
Select list of the most differentially expressed genes during co-culture with *L. crispatus* relative to *E. faecalis* single-species control.

| Old locus tag | Gene ID | Description | Log<sub>2</sub>FC | P-value | FDR |
| --- | --- | --- | --- | --- | --- |
| **Sugar and carbon utilization/uptake** |  |  |  |  |  |
| OG1RF_10030 |  | ABC superfamily ATP binding cassette transporter permease subunit | -2.6351 | 5.32E-05 | 1.82E-04 |
| OG1RF_10684 |  | maltose PTS enzyme II | 2.2417 | 6.60E-08 | 1.00E-06 |
| OG1RF_10991 |  | spermidine/putrescine ABC transporter permease protein | 3.2017 | 2.08E-05 | 8.18E-05 |
| OG1RF_10992 |  | spermidine/putrescine ABC transporter ATP-binding protein | 3.7275 | 2.59E-05 | 9.94E-05 |
| OG1RF_10993 |  | spermidine/putrescine ABC transporter substrate-binding protein | 3.9476 | 3.96E-06 | 2.14E-05 |
| OG1RF_10994 | ade | adenine deaminase | 2.4560 | 1.26E-04 | 3.75E-04 |
| OG1RF_11022 | trpX | tryptophan ABC transporter substrate-binding protein | 2.8249 | 3.55E-12 | 1.48E-09 |
| OG1RF_11023 |  | tryptophan ABC transporter permease protein | 2.2970 | 1.05E-10 | 1.75E-08 |
| OG1RF_11024 |  | tryptophan ABC transporter ATP-binding protein | 3.0645 | 3.95E-11 | 8.23E-09 |
| OG1RF_11133 | mdxG | sugar ABC transporter permease | 2.3167 | 1.72E-04 | 4.84E-04 |
| OG1RF_11134 | mdxF | sugar ABC transporter permease | 2.3274 | 7.82E-11 | 1.40E-08 |
| OG1RF_11135 | mdxE | extracellular maltose-binding protein | 3.0676 | 4.60E-13 | 2.92E-10 |
| OG1RF_11136 |  | glycoside hydrolase family 13 protein | 2.5327 | 9.58E-10 | 6.66E-08 |
| OG1RF_11137 |  | alpha-glucosidase | 2.2435 | 1.44E-08 | 3.56E-07 |
| OG1RF_11234 |  | PTS enzyme IIB component | 2.9516 | 5.79E-04 | 1.40E-03 |
| OG1RF_11542 |  | putative group I ECF transporter transmembrane (T) component | -2.0465 | 7.37E-09 | 2.38E-07 |
| OG1RF_11598 |  | PTS enzyme IIBC component | 2.5367 | 2.92E-09 | 1.38E-07 |
| OG1RF_11753 | treP | putative trehalose-specific PTS enzyme EIIBC component | 2.7823 | 5.03E-12 | 1.80E-09 |
| OG1RF_11763 |  | ABC transporter permease | -2.2422 | 6.13E-05 | 2.05E-04 |
| OG1RF_12087 |  | D-aminopeptidase superfamily protein | 2.2514 | 4.84E-07 | 4.23E-06 |
| OG1RF_12088 |  | putative glutamyl aminopeptidase | 2.7273 | 1.42E-08 | 3.55E-07 |
| OG1RF_12089 |  | oligopeptide ABC superfamily ATP binding cassette transporter, binding protein | 3.0122 | 1.47E-09 | 9.00E-08 |
| **Transcriptional regulators** |  |  |  |  |  |
| OG1RF_12163 | vanR | VanR-ABDEGLN family response regulator transcription factor | 2.5336 | 1.78E-07 | 2.04E-06 |
| OG1RF_12443 |  | Rrf2 family transcriptional regulator | 3.6850 | 3.19E-13 | 2.92E-10 |
| OG1RF_12573 | citO | DNA binding transcriptional regulator | -2.0584 | 1.58E-06 | 1.06E-05 |
| **Glycolipid modification** |  |  |  |  |  |
| OG1RF_10094 | sdaA | L-serine ammonia-lyase, alpha subunit | 2.1115 | 6.47E-09 | 2.35E-07 |
| OG1RF_10095 | serS | Serine tRNA ligase | 2.0975 | 3.72E-08 | 6.91E-07 |
| OG1RF_10030 |  | tRNA-Ala | -2.6351 | 5.32E-05 | 1.82E-04 |
| OG1RF_10328 |  | LysM domain-containing protein | 3.0385 | 3.24E-04 | 8.38E-04 |
| OG1RF_12461 | lrgA | antiholin-like murein hydrolase modulator | 2.3980 | 2.64E-03 | 5.43E-03 |
| OG1RF_12506 |  | LPXTG cell wall anchor domain-containing protein | 2.8139 | 8.31E-12 | 2.31E-09 |
| **Metabolism and stress response** |  |  |  |  |  |
| OG1RF_10288 | nhaC | putative transporter, Na<sup>+</sup>:H<sup>+</sup> antiporter family | -2.5366 | 7.05E-12 | 2.21E-09 |
| OG1RF_10597 | gloA4 | lactoylglutathione lyase | 2.1406 | 2.57E-08 | 5.26E-07 |
| OG1RF_10837 |  | AAA family ATPase | -2.1085 | 7.13E-09 | 2.38E-07 |
| OG1RF_11349 | pduQ | 1-propanol dehydrogenase PduQ | 2.6438 | 2.42E-04 | 6.50E-04 |
| OG1RF_11665 | cydB | cytochrome bd-I demethylquinol:oxygen oxidoreductase subunit II | -2.0955 | 3.47E-11 | 8.12E-09 |
| OG1RF_12165 | aadK | aminoglycoside 6-adenylyltransferase | 2.5423 | 1.25E-10 | 1.95E-08 |
| OG1RF_12442 |  | NAD(P)-dependent oxidoreductase | 3.8137 | 4.14E-14 | 1.04E-10 |
| OG1RF_12545 | rsmG | 16S rRNA (guanine(527)-N(7))-methyltransferase RsmG | 2.0277 | 3.49E-10 | 3.80E-08 |
| **Unknown function** |  |  |  |  |  |
| OG1RF_10023 |  | DUF308 domain-containing protein | -2.4806 | 6.75E-05 | 2.23E-04 |
| OG1RF_10264 |  | DUF4822 domain-containing protein | -2.4556 | 7.27E-09 | 2.38E-07 |
| OG1RF_10506 |  | Hypothetical protein | -2.5353 | 3.08E-10 | 3.67E-08 |
| OG1RF_10950 |  | Hypothetical protein | 2.0118 | 9.19E-06 | 4.16E-05 |
| OG1RF_11127 |  | Hypothetical protein | -2.6977 | 7.43E-05 | 2.41E-04 |
| OG1RF_11263 |  | Helix-turn-helix domain-containing protein | 2.0430 | 3.69E-10 | 3.85E-08 |
| OG1RF_11399 |  | Hypothetical protein | 3.0745 | 4.67E-13 | 2.92E-10 |

|  |  |  |  |  |
| --- | --- | --- | --- | --- |
| OG1RF_11794 | DoxX family protein | 4.7674 | 7.27E-13 | 3.64E-10 |
| OG1RF_12414 | LXG toxin | 2.2947 | 2.01E-02 | 3.33E-02 |
| OG1RF_12455 | TIGR00282 family metallophosphoesterase | 2.0867 | 7.09E-05 | 2.32E-04 |
| OG1RF_11937 | Hypothetical protein | 2.7725 | 5.05E-11 | 9.73E-09 |

Among the 20 highly up-regulated genes were the response regulator *vanR* (*OG1RF_12163*), two hypothetical proteins (*OG1RF_11794, OG1RF_11399*), putative LysM domain-containing peptidoglycan hydrolase (*OG1RF_10328*), FMN reductase (*OG1RF_12442*), Rrf2 family transcriptional regulator (*OG1RF_12443*), and spermidine/putrescine ABC family transporter (*OG1RF_10991-10993*), while the remaining genes encodes for proteins involved in central metabolism. Amongst the 20 most down regulated genes were malate quinone oxidoreductase (*OG1RF_10264*), cytochrome bd-l oxidoreductase (*OG1RF_11665*), four hypothetical proteins (*OG1RF_11127, OG1RF_10506, OG1RF_10023, OG1RF_10263*), an alanine tRNA ligase (*OG1RF_t10030*), and a putative Na^+^:H^+^ antiporter (NhaC; *OG1RF_10288*), while the remaining genes encode for proteins involved in sugar and carbon transport (**Fig 4A**).

### Transposon sequencing (Tn-seq) identifies genes that enable *E. faecalis* to compete with *L. crispatus*

Next, we used transposon sequencing (Tn-seq) ^67^ to identify *E. faecalis* OG1RF transposon mutants that are either under-represented (sensitivity to killing) or over-represented (resistant to killing) when co-cultured with *L. crispatus* . Using a false discovery rate (FDR) < 0.05 and Monte Carlos value of 0.373, eight mutants were under-represented at 24 hrs. Remarkably, all mutants contained insertions within the *dltA*, *dltB* or *dltD* genes (**Table S3**). The *dltABCD* operon (*OG1RF_12109-12112*) is responsible for D-alanylation of cell wall teichoic acid, a modification that is important for virulence and antimicrobial peptide and daptomycin tolerance in Gram-positive bacteria ^68–74^. Using the same cutoff applied to the 24 hrs time point, 30 Tn mutants were identified at 48 hrs, with three Tn mutants under-represented (*OG1RF_10193, OG1RF_10681* and *hexB*) and the remaining Tn mutants over-represented (**Table S4**). Among the over-represented genes with multiple hits, five had insertions within *ldh1* (*OG1RF_10199*), coding for the lactate dehydrogenase, two within *ndh2* (*OG1RF_11660*) coding for NADH dehydrogenase, and 12 within the *dltABCD* operon. At 72 hrs post incubation, 6 Tn mutants identified were under-represented, with 2 hits in genes of unknown function (*OG1RF_10435* and *OG1RF_11674*) and the remaining with disruptions in genes that contribute to different metabolic processes (**Table S5**).

Because insertions within the *dltABCD* operon and *ldh1* represented the biggest pool of Tn mutants identified, we assessed the *E. faecalis* arrayed transposon library ^75^ to retrieve individual *dltA* and *ldh1* Tn mutants in addition to three other Tn mutants (OG1RF_11697, OG1RF_10490 and *hexB*) that appeared at the 48 hrs and 72 hrs time point (**Table 2**) for validation purposes. While it is not a surprise that population of Tn mutants flux temporally when grown as a pooled library (in Tn-Seq), for example *dlt* Tn mutants being under-represented at 24 hrs and over-represented at 48 hrs, by individually co-culturing these Tn mutants with *L. crispatus*, we verified that the *dltA::*TnMar was more susceptible to *L. crispatus* killing, showing ∼1-log reduction in CFU recovered after 72 hrs when compared to the survival of the parent strain (**Fig 5A**). In agreement with the TnSeq hits, the *OG1RF_11697*::TnMar and *ldh1*::TnMar were, respectively, more sensitive (6-log) and more resistant (∼1-log) to *L. crispatus* killing. While statistically significant, Tn mutants with disruption in *OG1RF_12434* and *OG1RF_12308* were not drastically susceptible (∼1-fold reduction) to *L. crispatus*-mediated killing (**Fig 5A**). Again, 72 hrs viability of *L. crispatus* remained unchanged when co-cultured with any of these mutants (**Fig 5B**).

**Fig 5.**
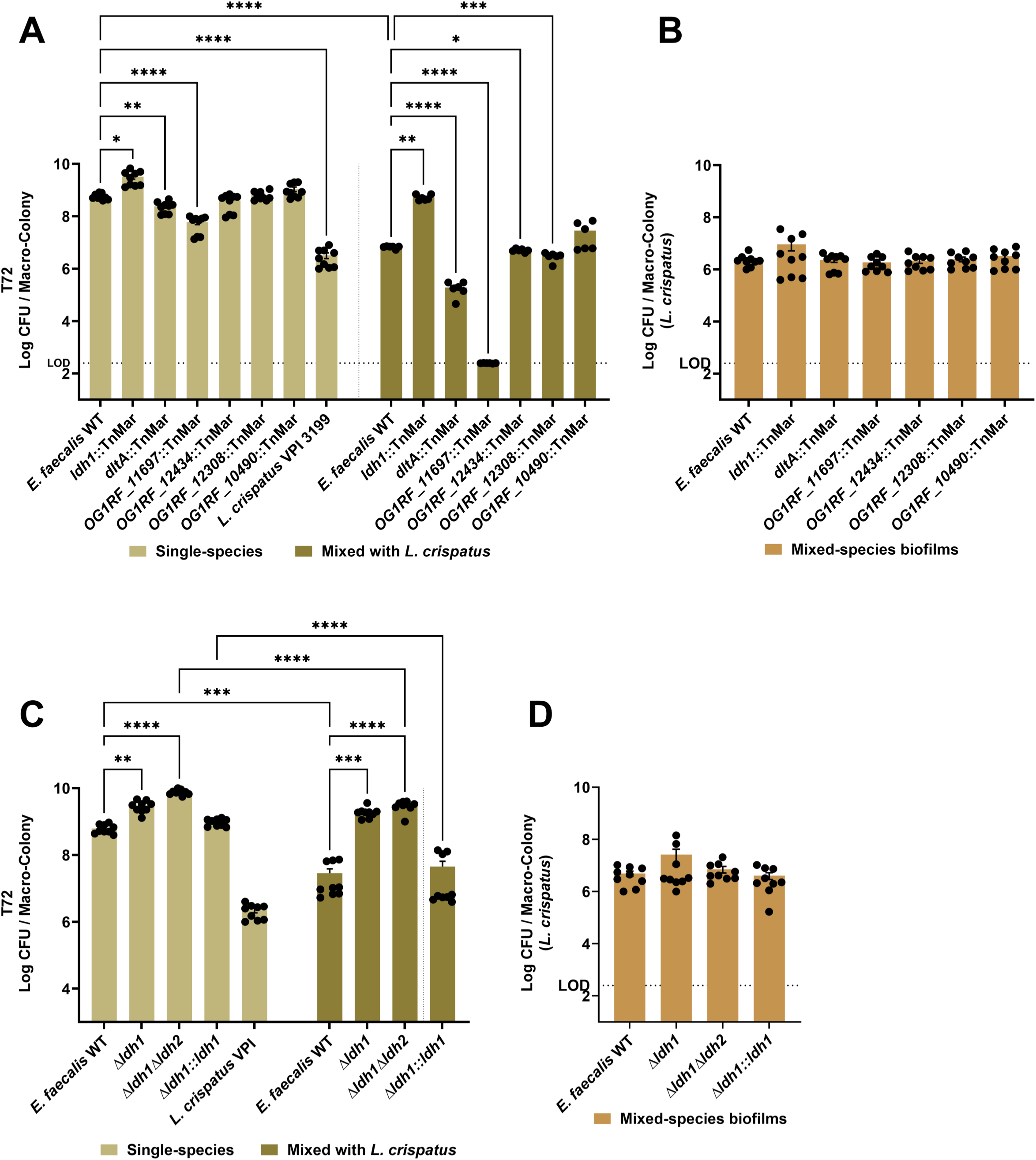
*E. faecalis* genes contribute to tolerance against *L. crispatus* antagonism. Colony-forming units (CFU) recovered from *E. faecalis* OG1RF, its isogenic transposon mutants, and *L. crispatus* VPI 3199 grown respectively either as single-species macro-colony biofilm, or as dual-species (*E. faecalis* + *L. crispatus*) macro-colony biofilm (A-B) for 72 hrs. Colony-forming units (CFU) recovered from *E. faecalis* OG1RF, its isogenic LDH deletion and complementation mutants, as well as *L. crispatus* VPI 3199 grown respectively either as single-species macro-colony biofilm, or as dual-species (*E. faecalis* + *L. crispatus*) macro-colony biofilm (C-D) for 72 hrs. Data points represent an average of 9-12 biological replicates, collated from at least three repeated experiments. Statistical analysis was performed using Brown-Forsythe ANOVA test with Welch’s correction. Error bars represent the standard error of margin (SEM). * *p* ≤ 0.05, ** *p* ≤ 0.01, *** *p* ≤ 0.001, **** *p* ≤ 0.0001.

**Table 2.** Selected list of differentially abundant *E. faecalis* transposon mutants during co-culture with *L. crispatus* relative to *E. faecalis*.

| Old locus tag | Gene Name | NCBI description | Single vs mixed t24 (Log<sub>2</sub>FC) | Single vs mixed t48 (Log<sub>2</sub>FC) | Single vs mixed t72 (Log<sub>2</sub>FC) |
| --- | --- | --- | --- | --- | --- |
| OG1RF_12109 | dltD | D-alanyl-lipoteichoic acid biosynthesis protein | -1.21 | 1.72 | ns |
| OG1RF_12111 | dltB | D-alanine transfer protein | -1.60 | 1.76 | ns |
| OG1RF_12112 | dltA | D-alanine--D-alanyl carrier protein ligase | -1.46 | 1.29 | ns |
| OG1RF_10199 | ldh1 | L-lactate dehydrogenase | ns | 2.42 | ns |
| OG1RF_11697 |  | Hypothetical protein | ns | 5.28 | ns |
| OG1RF_12434 | hexB | DNA mismatch repair protein | ns | -1.26 | ns |
| OG1RF_12435 |  | Hypothetical protein | ns | ns | -1.38 |
| OG1RF_12308 |  | Thioredoxin family protein | ns | ns | -4.21 |
| OG1RF_12309 | pepA2 | glutamyl aminopeptidase | ns | ns | -0.94 |
| OG1RF_10490 |  | Cell wall surface anchor family protein | ns | 3.06 | -4.21 |

In previous studies, *E. faecalis* LDH (known as Ldh1) was found to be important for iron-augmented energy production^76^ and for the inhibition of *P. aeruginosa* growth via L-lactate-dependent iron chelation^36^. In addition, a second LDH gene (*OG1RF_10373, ldh2*) is present in the OG1RF genome, sharing 46% similarity with Ldh1 ^36^. Even though Ldh1 has been found to play a prominent role in lactate production, both Ldh1 and Ldh2 have redundant activities participating in L-lactate production and redox maintenance ^36,77–79^. As further validation of the Tn screen, we tested the capacity of in-frame deletion strains lacking one or both *ldh* genes (Δ*ldh1*, *Δldh1Δldh2*) as well as a genetically-complemented *Δldh1* strain (*Δldh1::ldh1*) ^36^ to compete with *L. crispatus* using the macro-colony biofilm plate assay. As previously reported ^36^, inactivation of *ldh1* alone or in combination with *ldh2* enabled *E. faecalis* to grow to higher cell density (**Fig 5C**). Consistent with the TnSeq validation results, loss of *ldh1* alone, or both *ldh1* and *ldh2*, restored *E. faecalis* viability in the *L. crispatus*-*E. faecalis* macro-colony assay to levels observed in the single *E. faecalis* macro-colonies. Further, genetic complementation of *ldh1* (*Δldh1::ldh1*) restored sensitivity to killing as compared to survival rate of the parent strain (**Fig 5C**). Again, 72 hrs viability of *L. crispatus* remained unchanged when co-cultured with any of these mutants (**Fig 5D**).

Because the prevalence of lactobacilli are linked to urogenital and gastrointestinal microbiome homeostasis ^39–41,80^, and these microbes have been used as probiotics ^39–43,80–84^, in the next set of experiments, we explored the usefulness of the *Galleria mellonella* larvae model to examine the potential of *L. crispatus* to protect the *Galleria* larvae against *E. faecalis* lethal infection. First, we showed that *L. crispatus* does not kill *G. mellonella* as larvae inoculated with *L. crispatus* alone displayed similar survival rates as larvae injected with heat-killed bacteria (**Fig 6A**). Remarkably, co-inoculation of *L. crispatus* with *E. faecalis* enhanced larvae survival by ∼40% when compared to larvae inoculated with an identical dose of *E. faecalis* alone (**Fig 6B**). Next, we tested the virulence potential of the Δ*ldh1,* Δ*ldh2* and Δ*ldh1*Δ*ldh2* strains alone or co-inoculated with *L. crispatus.* While not statistically significant, all *ldh* mutants killed *G. mellonella* faster than parent or Δ*ldh1*::*ldh1* strains (**Fig 6C**). More importantly, the protection conferred by *L. crispatus* was completely lost in larvae co-infected with *L. crispatus* and any one of the *ldh* mutants (**Fig 6D**).

**Fig 6.**
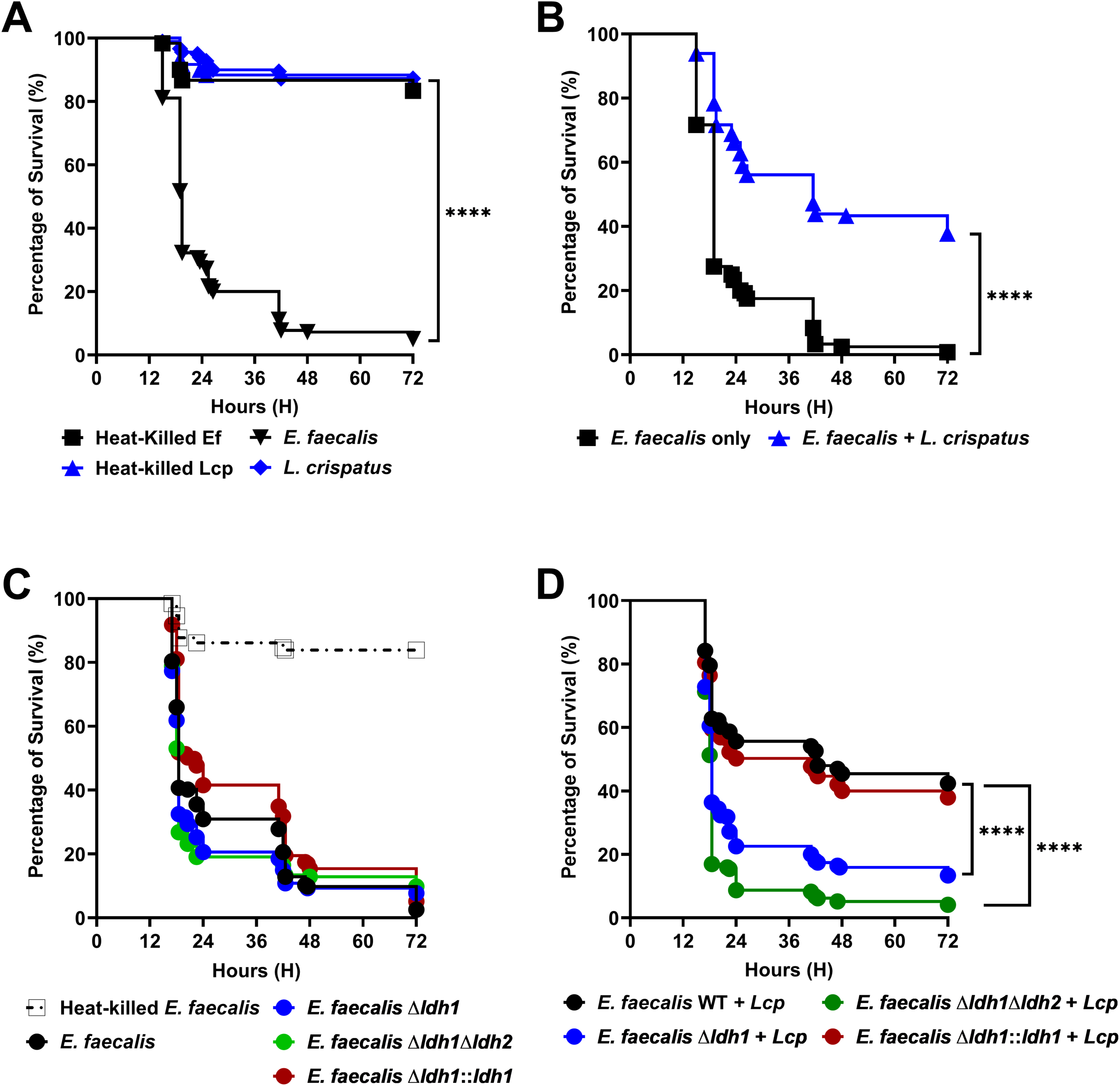
Loss of *ldh1* restores virulence of *E. faecalis* in a co-infection model of Galleria larvae. Percentage survival of *G. mellonella* larvae 72 hrs post-injection with either single-species controls; *E. faecalis* OG1RF or *L. crispatus,* and their heat-killed counterparts (A), or co-injected with both (B). Percentage survival of *G. mellonella* larvae 72 hrs post-injection with *E. faecalis* OG1RF parent strain and its isogenic LDH deletion and complementation strains alone (C), or their corresponding co-injected counterparts (D). Each curve represents a group of 20 larvae injected with 10^5^ CFUs suspended in PBS at a volume of 5 µL. Data points represent an average of 9 biological replicates. Statistical analysis was performed using Log-rank (Mantel-Cox) test. **** *p* ≤ 0.0001.

## DISCUSSION

Fundamental knowledge of microbial interactions is essential and has strong implications for the design of interventions for biofilm-associated infections ^18,85,86^. While there are reports implicating *E. faecalis* as a vaginal pathogen, the pathophysiology of *E. faecalis* in the vaginal tract is unknown. Since *L. crispatus* is a biomarker of vaginal health, and *E. faecalis* must outcompete vaginal lactobacilli to drive disease onset ^51–53,87^, by leveraging high throughput molecular approaches to identify genetic factors that contributes to competitive interactions between *E. faecalis* and *L. crispatus*, we will gain a better understanding of how *E. faecalis* mitigate assaults by *L. crispatus*.

Several studies have shown that cell-free supernatants of *L. crispatus* inhibits the growth of vaginal pathogens, including *Gardnerealla vaginalis* ^88^, *Candida albicans* ^89^ and *Streptococcus agalactiae* ^90,91^. In *S. agalactiae,* a major pathogen of vaginal infections, this inhibitory effect was largely attributed to the lower tolerance of *S. agalactiae* to the more acidic pH environment generated by the presence lactobacilli ^91^. Though our results did not completely concur with findings from other groups, that cell-free supernatants indeed have growth-inhibitory properties, it is certain that *L. crispatus* uses a contact-independent mechanism to kill *E. faecalis.* Perhaps, the molecule(s) responsible for killing *E. faecalis* might be labile; or alternatively, at inadequate dose in the supernatants. It is also plausible that acidic pH, phenyl-lactic acid (PLA) ^62^, and other complementary mechanisms that we are unaware of, work in concerted effort with these unknown molecule(s) to kill *E. faecalis*. Beside using antimicrobial molecules to curb outgrowth of vaginal pathogens, one recent study also noted that co-culturing with *L. crispatus* reduces *S. agalactiae* recto-vaginal colonization and attenuate invasion of endometrium decidual cells ^92^, though mechanisms remained to be determined, providing alternative explanations for *L. crispatus* ability to dominate the vaginal tract. Of similar context, a recent report also revealed that *L. crispatus* actively outcompetes *E. faecalis* to adhere to vaginal epithelial cells ^93^. While studies of *L. crispatus* are limited, *L. rhamnosus* that primarily cohabits the gastrointestinal tract has been extensively studied for its antagonistic capabilities ^80,84^ against other resident members of the gastrointestinal tract. Of these studies, the antagonistic mechanisms are speculated to be linked to secretion of bacteriocins, organic acids, or hydrogen peroxide, which are traits shared amongst members of the genus *Lactobacillus* ^49,66,84^. Concurring with previous studies that probe the potential of lactobacilli as probiotics ^42,43,82,83^, we also demonstrated that co-injection of *L. crispatus* with *E. faecalis* significantly enhanced *G. mellonella* larvae survival, and this *L. crispatus-*mediated protection was lost when larvae were co-injected with *L. crispatus* and isogenic *E. faecalis* LDH mutants that were resistant to *L. crispatus*-mediated killing. While results are promising, whether phenotypes will be recapitulated in a vaginal infection model remains to be addressed and will be pursued in the future. Though this report is focusing on how *E. faecalis* perceives the presence and attempt to compete with *L. crispatus*, studies to identify the molecular factors responsible for the killing of *E. faecalis* by *L. crispatus* will soon be underway.

By leveraging the killing ability of *L. crispatus* as a tool for Tn-seq screen, we identified *E. faecalis* genes that are important for competition against *L. crispatus*. We found that *E. faecalis* Tn mutants with disruption in the *dltABCD* operon or *OG1RF_11697* (hypothetical protein) were more susceptible to *L. crispatus* killing, whereas Tn mutants with disruption in the *ldh1* gene showed increased resistance. The *dltABCD* gene products that catalyze the addition of d-alanine esters to wall teichoic acids ^94^ and lipoteichoic acids (LTA) ^95^, two important components of the cell wall of gram-positive bacteria, are essential in the regulation of autolysis, cation homeostasis, host cell adhesion and invasion, and biofilm formation, and is essential for virulence of pathogens such as *Listeria monocytogenes* and *S. aureus*, including *E. faecalis* ^95^. Most importantly, the D-alanylation of teichoic acids provides resistance to cationic antimicrobial peptides (CAMP), by raising the overall net negative charge of the membrane, thereby reducing the affinity for CAMP^96^. Since the loss of *dltA* in *E. faecalis* that encodes for a D-alanyl protein ligase, enhances sensitivity to CAMP ^97,98^, it is tempting to speculate that *L. crispatus* produces antimicrobial peptide(s) that kill *E. faecalis*.

In agreement with our speculation, we observed an up-regulation of the *vanRS* two component system, including *vanY_B_* that codes for an orphan putative serine-type D-Ala-D-Ala carboxypeptidase (Table 1, Table S1) in the RNAseq. In enterococci, the VanRS two component system (TCS) regulate a downstream vancomycin (Van) resistance accessory operon that is responsible for the modification of the terminal d-alanyl-d-alanine (d-Ala-d-Ala) dipeptide of the muramyl pentapeptide ^99^, a precursor in peptidoglycan synthesis, and in doing so, prevent the binding of Van and cell death. Specifically, by coupling with the appropriate protein ligase, the incorporation of either terminal d-Ala-d-Lac or d-Ala-d-Ser dipeptides, prevent Van-mediated inhibition of peptidoglycan cross-linking ^100^. If indeed an antimicrobial peptide is facilitating *L. crispatus* killing, we anticipate resistant to killing in a VanRS-responsive manner. Ironically, apart from an orphan VanY encoded downstream of VanRS, *E. faecalis* OG1RF genome lacks a complete Van accessory operon. Paradoxically, *E. faecalis* V583 which is more susceptible to *L. crispatus* mediated killing than OG1RF, possesses a Van accessory operon downstream of its VanRS_B_ (type B) TCS (*EF_2299-2298;* 32% and 22.4% a.a identity respectively to VanRS of OG1RF). Like V583^101^, another Van resistant enterococcal strain, *E. faecalis* ATCC 51299 increased synthesis of d-Ala-d-Lac-terminated peptidoglycan pentapeptides to prevent Van-mediated inhibition of peptidoglycan cross-linking ^102^. Perhaps, having more d-Ala-d-Lac-terminated pentapeptides render *E. faecalis* V583 more vulnerable to *L crispatus* mediated killing. Besides the fact that a Zn-dependent D, D-carboxypeptidase (*OG1RF_12161*; 52% a.a) (annotated as *vanY* in V583 genome; for the cleavage of D-Ala-d-Ala dipeptides) is of close proximity to VanRS in the core genome, studies of VanRS-facilitated remodeling of peptidoglycan in OG1RF has not been pursued. Ultimately, what we gathered from these findings is that the enterococcal cell wall interface seems to be the target of *L. crispatus* killing. Further studies are warranted to discern the underlying mechanisms.

Besides *dltABCD* operon, *OG1RF_11697* was found to greatly contribute to *E. faecalis* fitness. Notably, *OG1RF_11697* codes for a hypothetical protein homologous to *L. monocytogenes* (37% similarity) and *S. aureus* (97% similarity) YbbR proteins. Like *L. monocytogenes* and *S. aureus*, *OG1RF_11697* (henceforth *ybbR*) resides in a conserved three-gene operon that includes the diadenylate cyclase CdaA (*OG1RF_11698*), and phosphoglucosamine mutase GlmM (*OG1RF_11696*). GlmM which is required for production of glucosamine-1-phosphate, an early intermediate of peptidoglycan synthesis and CdaA is the sole enzyme responsible for c-d-AMP synthesis in *E. faecalis* ^103^. While mechanisms remain poorly defined, recent studies have shown that YbbR and GlmM fine-tune intracellular c-di-AMP levels by enhancing or inhibiting the ability of CdaA to synthesize c-di-AMP. Specifically, co-expression of *S. aureus* CdaA with YbbR increased intracellular c-di-AMP pools whereas co-expression of CdaA with GlmM reduced c-di-AMP levels ^104^. In *Lactococcus lactis*, a single amino acid substitution in GlmM resulted in a tighter interaction with CdaA and, as a result, triggered a drastic decrease in intracellular c-di-AMP levels ^105^. C-di-AMP regulates a wide-range of cellular processes, including but not limited to osmoregulation, central metabolism, and virulence ^106,107^. Because of the strong conservation of the *cdaA-ybbR-glmM* operon in Gram positive bacteria, and multiple evidence associating YbbR and GlmM with c-di-AMP production, we speculate that the increased sensitivity of the *ybbR*::TnMar mutant to *L. crispatus* killing is linked to c-di-AMP production.

Unlike the decreased fitness observed in *dltABCD* and *ybbR* mutants, an explanation for the increased survival of *ldh1* mutants is less straightforward. One possibility is that the profound alteration in the fermentation end-product profile of Δ*ldh1* strains, due to funneling of pyruvate to other enzymes in glycolysis, changing from mainly homolactic to heterolactic i.e., a distinct characteristic of the LDH-null strain that interferes with *L. crispatus* ability to produce anti-*E. faecalis* metabolites. Specifically, an *E. faecalis Δldh1Δldh2* mutant that produces negligible amounts of lactate displays significant increases in formate (∼4-fold), ethanol (∼3-fold), and acetoin (∼5-fold) produced ^108^. While the mechanism is unknown, both formate and ethanol have been shown to modulate growth of lactobacilli. Specifically, co-culturing lactobacilli with *Saccharomyces cerevisiae* enhanced lactic acid production by lactobacilli while decreasing ethanol production by *S. cerevisiae* ^109^. In another example, formate produced by *Streptococcus thermophilus* stimulated *L. delbrueckii* growth and exopolysaccharide (EPS) production ^110^. Another possible, non-mutually exclusive explanation is that the metabolic alterations associated with the loss of LDH activity enhance *E. faecalis* overall fitness.

In summary, the findings presented here represent an important first step towards the identification of genetic determinants that allow *E. faecalis* to mitigate *L. crispatus* killing, which may have far-reaching implications to other ecological niches, i.e. gastrointestinal tract, where members of both genera co-exist. In the long-run, findings here will provide explanations for the outgrowth of *E. faecalis* in the vaginal tract, and consequently, vaginal infection.

## MATERIAL & METHODS

### Bacterial strains and growth conditions

The bacterial strains used in this study are listed in **Table 3**. Bacteria were grown in either brain heart infusion (BHI) broth (for enterococci) or de Man, Rogosa and Sharpe (MRS) broth (lactobacilli) (both from BD Difco, New Jersey, USA) at 37°C under microaerophilic conditions (5% CO_2_). For growth kinetic assays, overnight cultures were adjusted/normalized to an optical density (OD_600_) of 0.25 (for enterococci) or 0.75 (for lactobacilli) (∼1 × 10^8^ colony forming units (CFU) mL^-1^) and inoculated into fresh media at 1:50 ratio, with changes in OD_600_ over time recorded in an automated growth reader (Bioscreen c, Oy Growth Curves AB, Helsinki, Finland). Unless stated otherwise, starter cultures for *in vitro* and *in vivo* experiments were adjusted from overnight cultures to the desired OD_600_ using phosphate-buffered saline (PBS). All biofilms were grown at 37°C under microaerophilic conditions (5% CO_2_) in MRS broth. For *E. faecalis* OG1RF CFU determination, serially diluted aliquots were plated on BHI agar supplemented with 200 µg mL^-1^ rifampicin and 10 µg mL^-1^ fusidic acid. For CFU determination of other enterococci, bile aesculin azide (BAA) agar (BD Difco) was used. For lactobacilli CFU determination, serially diluted aliquots were plated on Rogosa agar (Oxoid, Massachusetts, USA). Chemicals and biological reagents were purchased from Sigma Aldrich (Missouri, USA) unless stated otherwise.

**Table 3.**
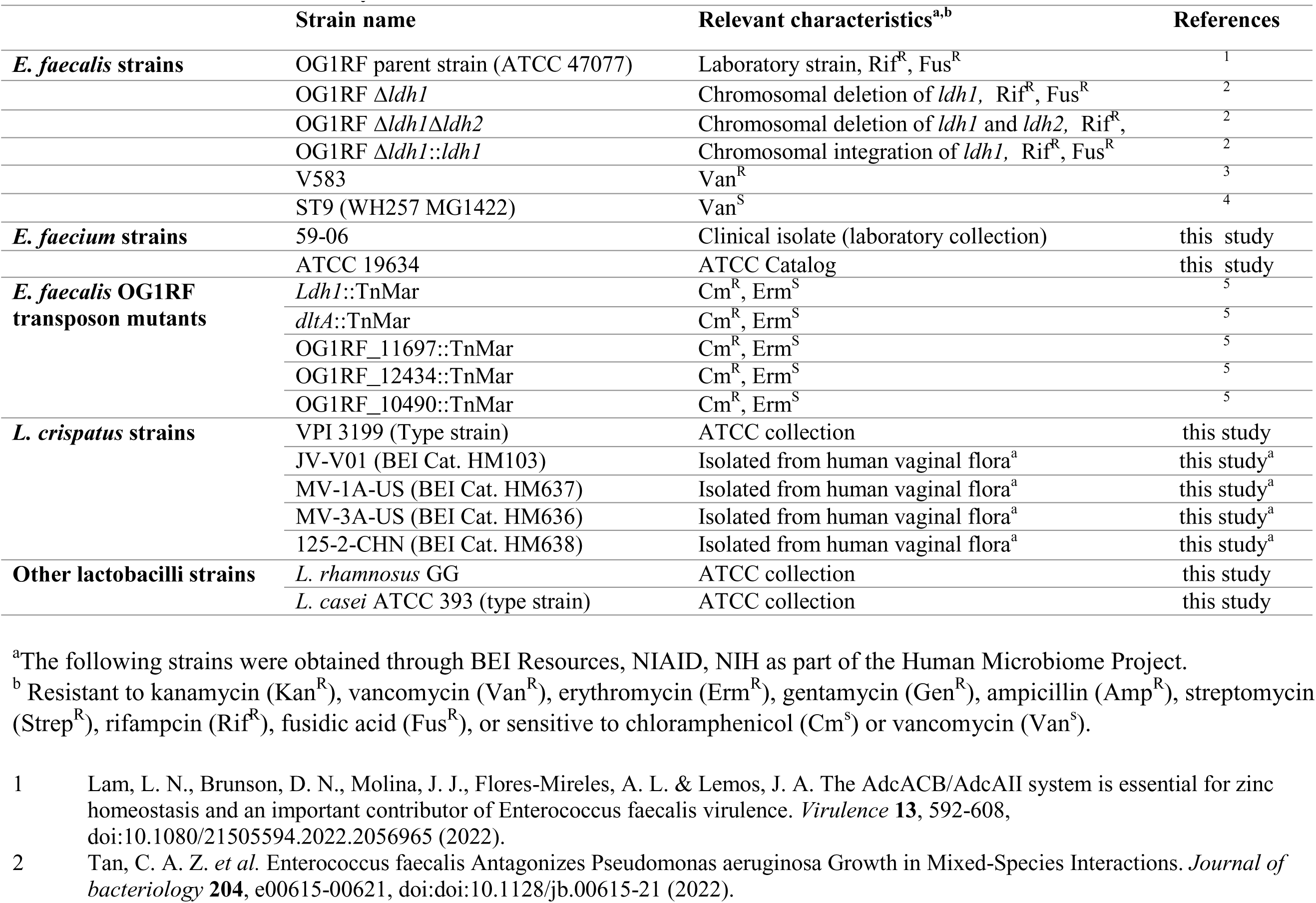
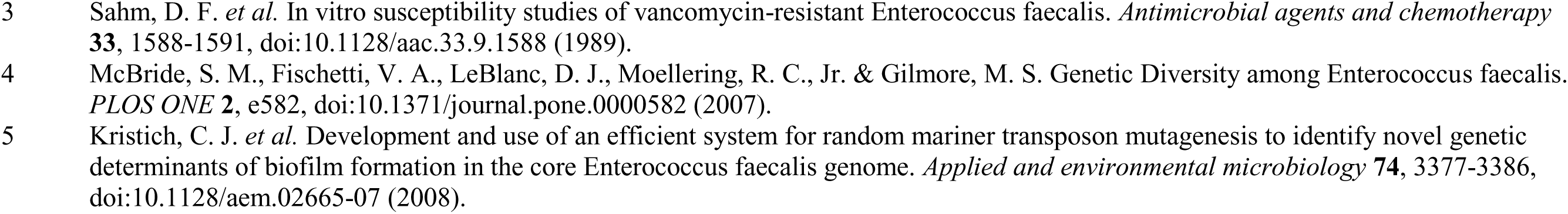
Strains used in this study.

### Biofilm assay

Biofilm assays were performed as previously described with minor modifications. For macro-colony biofilm, 1 × 10^6^ CFU of each bacterium in a total volume of 5 µL was inoculated onto the surface of MRS agar. After incubation at selected timepoints, macro-colonies were excised using a scalpel and homogenized by vortexing in PBS, followed by plating on selective media for CFU enumeration. For single and two-species biofilms established in broth, 1 × 10^6^ CFU of bacterial culture suspended in a total volume of 20 µL PBS was inoculated into 1 mL of MRS broth in a 24-well microtiter tissue culture plate. To investigate if *L. crispatus*-mediated killing was contact-dependent or independent, biofilms were allowed to form in a 6-well tissue culture plate with a Transwell™ insert (Corning, New York, USA). To investigate if *L. crispatus*-secreted molecules are responsible for *E. faecalis* killing, biofilm supernatants (either single or dual-species biofilms grown for 72 hrs) were recovered and filter sterilized using 0.2-micron polyether sulfone membrane filters (Millipore, Massachusetts, USA), then mixed at equal ratio (v/v) with MRS media to grow *E. faecalis* biofilms.

### Phenyl-lactic acid (PLA) susceptibility assay

Bacteria were grown overnight in MRS media, adjusted to an OD_600_ of 0.5 in PBS, and diluted at a ratio of 1:1000. Diluted cultures were inoculated at a ratio of 1:20 into MRS broth containing increasing concentrations of phenyl-lactic acid and incubated at 37°C with 5% CO_2_ for 24 hrs. The absorbance at OD_600_ was measured using Synergy H1 microplate reader (Molecular Devices, California, USA).

### Spot antagonism assay

Bacterial spot biofilms were prepared as previously described ^111^ with minor modifications. Briefly, 10 µL of normalized starter cultures were spotted on MRS plates with *E. faecalis* inoculated either at the same time or 24, 48, or 72 hrs after *L. crispatus*.

### Galleria mellonella infection

To assess *E. faecalis* virulence, the larvae of *G*. *mellonella* were infected as described previously ^112^. Briefly, larvae (groups of 20) were injected with either i) *E. faecalis* OG1RF or *L. crispatus* VPI 3199 alone (∼ 5 × 10^5^ CFU), ii) heat-killed OG1RF or VPI3199 (60 mins at 100°C), or iii) dual-species inoculum. Post-injection, larvae were kept at 37°C and their survival recorded over time for 72 hours.

### Transposon mutant library growth as dual species macro-colony biofilms

The *E. faecalis* OG1RF pooled Tn library ^67^ was recovered from glycerol stock and suspended in 1 mL of PBS. Single-species *E. faecalis* pooled Tn library and *L. crispatus* macro-colony biofilms were produced by inoculating 1 × 10^6^ CFU of each in a total volume of 5 µL onto the surface of MRS agar. To produce dual-species macro-colony biofilms, equal volume of single-species *E. faecalis* pooled transposon library and *L. crispatus* were mixed and inoculated onto the surface of MRS agar. After incubation at selected timepoints, macrocolonies were excised using a scalpel and suspended in PBS. For Tn-seq, single-species and dual-species macrocolony biofilm samples were prepared accordingly, then pelleted at 4,000 x rpm for 10 min, and frozen at -80°C until further use.

### Transposon library sequencing (Tn-seq) analysis

The Wizard DNA purification kit (Promega, Wisconsin, USA) was used to obtain genomic DNA from single-species or dual-species macro-colony pellets (prepared in triplicates). Samples were submitted to the University of Minnesota Genomics Center for library preparation and sequencing. Sequencing was performed using an Illumina NextSeq 2000 in 150-bp paired-end output mode. Tn-seq analysis was performed with custom scripts using the University of Florida supercomputer HiperGator. Analysis of sequencing reads were performed as previously described ^113^. Briefly, bioinformatics reads were mapped to the OG1RF reference genome (GenBank CP002621.1, NCBI RefSeq NC_017316.1), and Tn insertions at TA sites quantified. The statistical significance of the relative abundance of Tn reads at each TA site in each corresponding single-species biofilm compared to dual-species biofilm samples was evaluated using a chi-square test and a Monte Carlo-based method. Th scripts used in this study are publicly available (https://github.com/dunnylabumn/Ef_OG1RF_TnSeq). Log_2_ fold changes were calculated from each corresponding single-species biofilm compared to dual-species biofilm samples. Statistical significance was defined as a *P* value of <0.05 and a Monte Carlo simulation value of 3.73184 (the lowest value obtained in these calculations). Tn mutants used for additional experiments were obtained from frozen library stock plates (gifts from Dunny lab, University of Minnesota, USA). Verification of Tn mutants were performed as previously described using selective plating and sequencing ^75,114^.

### RNA sequencing and analysis

Triplicates of single-species *E. faecalis* static biofilms were produced by inoculating 1 × 10^6^ CFU in a total volume of 5 mL MRS media in a 6-well tissue culture plate. To produce co-culture biofilms, equal inocula of single-species *E. faecalis* and *L. crispatus* cells were inoculated into the same well of a tissue culture plate. Separated by a Transwell™ (Corning, New York, USA) permeable membrane insert that prevent physical, *E. faecalis* biofilms were seeded on the surface of the Transwell™ permeable membrane, whereas *L. crispatus* biofilms were grown on the bottom of the tissue culture plate. After incubation for 24 hrs, biofilm samples were pelleted at 4,000 x rpm for 10 min, then suspended in RNAprotect solution (Qiagen, Hilden, Germany) for 5 mins. Then, samples were pelleted at 4,000 x rpm for 10 min again for downstream RNA extraction. Subsequent RNA extraction steps were performed according to the manufacturer’s protocol using PureLink RNA mini kit (Thermofisher, Massachusetts, USA) with minor modifications. Briefly, samples were incubated with 30 mg ml^-1^ of lysozyme for 30 mins prior to extracting RNA. After extraction, RNA samples were further processed using Monarch RNA cleanup kit (New England Biolabs, Massachusetts, USA). Then, the extracted RNA was quality checked using the RNA ScreenTape on a TapeStation instrument (Agilent Technologies, USA). Every sample had a minimum RNA concentration of 25 ng/µL and a RINe value ≥8.0. Extracted RNA samples were sent to SeqCenter (Pennsylvania, USA) for library preparation, ribosomal RNA depletion, sequencing and downstream bioinformatic analysis. Library preparation was performed using Illumina’s Stranded Total RNA Prep Ligation with Ribo-Zero Plus kit and 10bp unique dual indices (UDI). Sequencing was done on a NovaSeq X Plus, producing paired end 150bp reads. Demultiplexing, quality control, and adapter trimming was performed with illumina software bcl-convert (v4.2.4). Read mapping was performed with HISAT2^115^. Read quantification was performed using Subread’s featureCounts^116^. Read counts were normalized using edgeR’s Trimmed Mean of M values (TMM) algorithm^117^. Subsequent values were then converted to counts per million (CPM). Differential expression analysis was performed using edgeR’s glmQLFTest. Differentially expressed genes were considered using a cut-off of log_2_ fold change (between ≤-1 and ≥1) and *p* ≤ 0.05. Finally, pathway analysis was performed using limma’s *“kegga”* functionality^118^. The genes that were considered Up/Down in this analysis were set at a cut-off using Benjamini-Hochberg adjusted *p* ≤ 0.05.

### Statistical analysis

Data obtained from this study were analysed using GraphPad Prism 9.0 software (GraphPad Software, San Diego, California, USA). Data from multiple experiments conducted on non-consecutive days were collated and applicable statistical tests were used.

## ACKNOWLEDGEMENT

We would also like to express our gratitude to Dr Julia Willet and Dr Gary Dunny for kindly providing the *E. faecalis* OG1RF transposon (SmarT) library and discussions of the Tn-seq results. This study was partially supported by the University of Florida College of Dentistry Seed grant (ID: 00132632) to LNL and UF Preeminence funds to JAL.

## DATA AVAILABILITY STATEMENT

The authors confirm that the data supporting the findings of this study are available within the article or in the supplementary material. The sequencing files generated from Tn-Seq (BioProject ID: PRJNA1170924) and RNA-seq (GEO Accession ID: GSE279275) have been deposited with the NCBI Sequence Read Archive and Gene Expression omnibus database.

---

**Fig S1. *L. crispatus* antagonism of enterococci is conserved at the species level.** Colony-forming units (CFU) recovered from *L. crispatus* clinical strains grown either as single-species macro-colony biofilm, or as dual-species (with *E. faecalis* OG1RF) macro-colony biofilm for 72 hrs (A). Colony-forming units (CFU) recovered from *L. crispatus* VPI 3199 grown either as single-species macro-colony biofilm, or as dual-species (with clinical enterococcal strains) macro-colony biofilm for 72 hrs (B). Data points represent an average of 9-12 biological replicates, collated from at least three repeated experiments. Statistical analysis was performed using Brown-Forsythe ANOVA test with Welch’s correction. Error bars represent the standard error of margin (SEM). * *p* ≤ 0.05, ** *p* ≤ 0.01, *** *p* ≤ 0.001.

**Fig S2. *L. rhamnosus* and *L. casei* are antagonists.** Colony-forming units (CFU) recovered from *E. faecalis* OG1RF, *L. casei*, and *L. rhamnosus* respectively when grown either as single-species macro-colony biofilm, or as dual-species macro-colony biofilms (with *L. casei* (A) or with *L. rhamnosus* (B)) at 24 and 72 hrs. Data points represent an average of 9-12 biological replicates, collated from at least three repeated experiments. Statistical analysis was performed using Brown-Forsythe ANOVA test with Welch’s correction. Error bars represent the standard error of margin (SEM). * *p* ≤ 0.05, ** *p* ≤ 0.01, *** *p* ≤ 0.001, **** *p* ≤ 0.0001.

**Fig S3. *L. crispatus* biofilm antagonistic activity is strengthened when in broth-based assays.** Colony-forming units (CFU) recovered from *E. faecalis* OG1RF (A) and *L. crispatus* VPI 3199 (B) when grown statically either as single-species biofilm, or as dual-species (with *E. faecalis* OG1RF) biofilm for up to 72 hrs using tissue culture plates. Data points represent an average of 9 biological replicates, collated from three repeated experiments. Statistical analysis was performed using Brown-Forsythe ANOVA test with Welch’s correction. Dotted line represents limit of detection; CFU of 42. Error bars represent the standard error of margin (SEM). **** *p* ≤ 0.0001.

**Fig S4. Lactobacilli and *E. faecalis* have similar tolerance to lactobacilli-derived phenyl-lactic acid *in vitro*.** Growth of *E. faecalis* OG1RF and *L. crispatus* VPI 3199 in MRS media supplemented exogenously with increasing concentration of phenyl-lactic acid (PLA) after 24 hrs. Data points present an average of 3 biological replicates. Statistical analysis was performed using Wilcoxon matched pairs signed rank test. Error bars represent standard deviations (S.D).

**Fig S5. Testing of biofilm supernatants for anti-enterococcal molecules by altering media ratios.** Colony-forming units (CFU) recovered from *E. faecalis* OG1RF growth after 24 hrs in MRS media mixed with 72 hrs cell-free biofilm supernatant isolated from single-species and dual-species biofilms, that was mixed at 90:10 ratio with fresh MRS, then adjusting media to final pH of 6.5. Data points represent an average of 9-12 biological replicates, collated from at least three repeated experiments. Statistical analysis was performed using Brown-Forsythe ANOVA test with Welch’s correction. Error bars represent the standard error of margin (SEM).

**Table S1. List of differentially expressed genes during co-culture with *L. crispatus* relative to *E. faecalis* single-species macro-colony biofilm.**

**Table S2. KEGG analysis of pathways differentially changed during co-culture relative to *E. faecalis* alone.**

**Table S3. List of differentially abundant *E. faecalis* transposon mutants during co-culture with *L. crispatus* relative to *E. faecalis* transposon library single-species macro-colony biofilm after 24 hr.**

**Table S4. List of differentially abundant *E. faecalis* transposon mutants during co-culture with *L. crispatus* relative to *E. faecalis* transposon library single-species macro-colony biofilm after 48 hr.**

**Table S5. List of differentially abundant *E. faecalis* transposon mutants during co-culture with *L. crispatus* relative to *E. faecalis* transposon library single-species macro-colony biofilm after 72 hr.**

